# Non-respiratory functions of *Saccharomyces cerevisiae* mitochondria are required for optimal attractiveness to *Drosophila melanogaster*

**DOI:** 10.1101/007336

**Authors:** Kelly M. Schiabor, Allison S. Quan, Michael B. Eisen

## Abstract

While screening a large collection of wild and laboratory yeast isolates for their ability to attract *Drosophila melanogaster* adults, we noticed a large difference in fly preference for two nearly isogenic strains of *Saccharomyces cerevisiae*, BY4741 and BY4742. Using standard genetic analyses, we tracked the preference difference to the lack of functional mitochondria the stock of BY4742 used in the initial experiment. We used gas chromatography coupled with mass spectroscopy to examine the volatile compounds produced by BY4741 and the mitochondria-deficient BY4742, and found they differed significantly. We found that several ethyl esters are present at much higher levels in strains with functional mitochondria, even in fermentative conditions. We confirmed the role of these ethyl esters in attraction by examining an EEB1Δ strain which reduces ethyl ester production. We found that nitrogen levels in the substrate affect the production of these compounds, and that they are produced at high levels by strains with functional mitochondria in the fermentation of natural substrates. Collectively these observations demonstrate the effect core metabolic processes have in mediating the interaction between yeasts and insect vectors, and highlight the importance of non-respirative mitochondrial functions in yeast ecology.

## Introduction

Since the early 20th century, the vinegar fly *Drosophila melanogaster* and the brewer's yeast *Saccharomyces cerevisiae*, have been the preeminent model species of molecular biology. Although generally studied by separate sets of researchers in the lab, these species are found together in nature on fermenting fruit substrates – an interaction that has received surprisingly little attention.

Their co-occurrence is no coincidence: each species benefits from the presence of the other. Ethanol produced by *S. cerevisiae* protects *D. melanogaster* larvae from predators [1], and *D. melanogaster* larvae preferentially feed on *S. cerevisiae*, which provides a complete nutritional source for larval development [2, 3]. The non-motile *S. cerevisiae*, in turn, rely on *D. melanogaster* and other insects to move to fresh, nutrient-rich substrates [4], and the burrowing activities of *D. melanogaster* larvae expose nutritional sources to the yeast that would have been otherwise inaccessible [5].

Given the benefits that each species gains from this interaction, it is not surprising that both yeast and fly appear to have evolved molecular mechanisms to actively maintain their co-localization.

It is well known to anyone who keeps fruits in their kitchen that *D. melanogaster* are attracted to fermenting fruit: an effect mediated almost entirely by volatile molecules produced by yeasts, and not the fruity substrate [2], although the substrate can have a significant effect on what compounds are produced[6–12]. Numerous volatile compounds produced by *S. cerevisiae* activate specific odorant receptor neurons in the *D. melanogaster* antennae, which are relayed to the odor representation center of the fly brain, the antennal lobe [13, 14], and elicit specific attractive and repulsive responses [2, 15–22].

The response to individual compounds is complex, and includes various types of attraction and avoidance behaviors. A small pulse of ethyl butyrate, for example, is sufficient to keep a fly flying towards the source of the odorant for twenty minutes without additional application of the odor, but this tracking is much less pronounced for acetic acid, another attractive yeast-produced volatile [17]. Mixtures of yeast-produced volatile compounds, more akin to what flies would experience in nature, produce more complex, context-dependent responses [16].

Different yeast species produce varying volatile bouquets, even when grown on identical substrates [6, 8, 12, 23–26]. This variance in volatile bouquets is observed across *S. cerevisiae* strains as well[24]. Further, fly species, and also different *D. melanogaster* strains, show varying preferences for yeast strains and species [27–29].

We became interested in the possibility that the large repertoire of genetic tools available for these species would allow us to exploit these different types of variation to better understand the molecular details of this interaction. As a first step, we examined the response of a single *D. melanogaster* strain to different *S. cerevisiae* strains grown under identical conditions.

## Results

### Two nearly isogeneic lab strains vary in their attractiveness to *D. melanogaster*

We used a simple preference assay (see Methods) to screen a large panel of *S. cerevisiae* strains, collected from diverse environments with varied histories of laboratory use and propagation, for their attractiveness to a wild-caught *D. melanogaster* line (Raleigh 437). We found considerable variation in the attractiveness across the strains (corroborating the results of [24]). As we were ultimately interested in exploring the genetic basis (in yeast) for this varied *D. melanogaster* attractiveness, we also included the two nearly isogeneic haploid *S. cerevisiae* strains, BY4741 and BY4742, which had been used to generate the systematic yeast deletion collection [30]. We were astonished to find that traps baited with BY4741, the MAT a parent, consistently attracted more flies when directly competing against traps baited with BY4742, the MAT*α* parent (Figure 1). The remainder of this paper is focused on understanding the origins of this preference difference.

**Figure 1.**
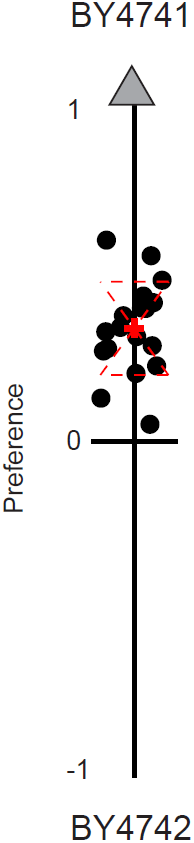
*D. melanogaster* are preferentially attracted to BY4741 over nearly isogenic BY4742. We compared *D. melanogaster* preference for different yeast strains using an open-flight preference assay. Each black dot represents the results of a single open-flight preference trial between lab stocks of BY4741 and BY4742. The preference index (PI) for each trial is defined as: PI = (A-B)/(A + B/2), where A = total flies caught in BY4741 baits and B = total flies caught in BY4742 baits. This value represents the direction of the preference (positive for A, negative for B) as well as the normalized degree of preference by the percentage over the expected. Red dashed lines represent the two-tailed standard deviation and the red “+” represents the mean of all trials.

### Attraction difference between BY4741 and BY4742 is associated with functional mitochondria

Outside of the mating type locus, these strains ostensibly differed at two metabolic marker loci: MET17 (deleted in BY4741) and LEU2 (deleted in BY4742). Given the known relationship between amino acid metabolism and volatile compound production [6, 9, 10, 31–35], we suspected that the difference in behavioral response to these two strains could be traced to their respective auxotrophies. We therefore crossed BY4741 and BY4742 and dissected several tetrads to obtain haploid strains with all eight possible genotypes at the three variable loci (MAT, LEU2 and MET15). However, when we tested these strains using our choice assay against BY4742, we found that all eight genotypes were strongly preferred to BY4742, including the MAT*α*; MET15; leu2Δ strain, which has the identical three-locus genotype as BY4742 (Figure 2A). We tested a subset of these strains against BY4741, and found that the flies preferred each strain equally (Figure 2B).

**Figure 2.**
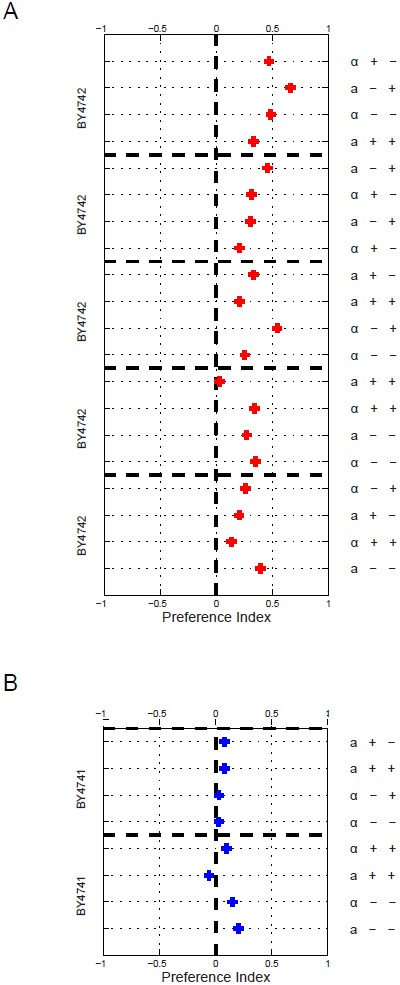
Genetic analysis of *D. melanogaster* preference for BY4741 over BY4742 suggests mitochondrial inheritance. To test if behavioral preference is associated with genotype, BY4741 and BY4742 were crossed and sporulated and the resulting haploid segregants genotyped at the MAT, MET15, and LYS2 loci. Segregants from five tetrads, collectively covering spores of every genotype, were tested against BY4742 in our behavioral assay. The red “+” represents the mean preference index of three trials. A) Tetrad spores of all genotypes phenocopy the BY4741 parent, suggesting a mitochondrial inheritance pattern. B) Two tetrads were tested against both BY4741 (blue “+”s), with the “+” representing the mean preference index of five behavioral trials. We found no preference between each tetrad spore and BY4741, all of which possess functional mitochondria. As shown in (A), all spores were preferred over BY4742, which does not possess functional mitochondria.

This non-autosomal inheritance pattern immediately suggested mitochondrial linkage. We tested both parental strains and all progeny for growth on glycerol, a non-fermentable carbon source, and found that all of the strains grew normally, except for BY4742 (Figure S1). This result confirmed a respiration deficiency in the BY4742 strain used in this study, which we refer to hereon as BY4742p, for petite.

To corroborate that the absence of a functional mitochondria reduces the attractiveness of yeast strains to *D. melanogaster*, we tested an independent BY4742 isolate (BY4742g, for grande), with a functional mitochondria (*MATα;his3;leu2;lys2;ura3*), as well the MRPL16Δ strain (*MATα;his3;leu2;lys2;ura3;mrpl16*) from the same deletion collection. MRPL16 encodes for the large subunit of the mitochondrial ribosomal protein, the deletion of which results in a respiration deficiency. We found that the independent BY4742g was equally as attractive as BY4741, while MRPL16Δ strain was less attractive than BY4741. Following the trend found with BY4741, we found BY4742g was also more attractive than both BY4742p and the MRPL16Δ strain (Figure 3). This further confirms that *D. melanogaster* are more attracted to S*. cerevisiae* strains possessing functional mitochondria.

**Figure 3.**
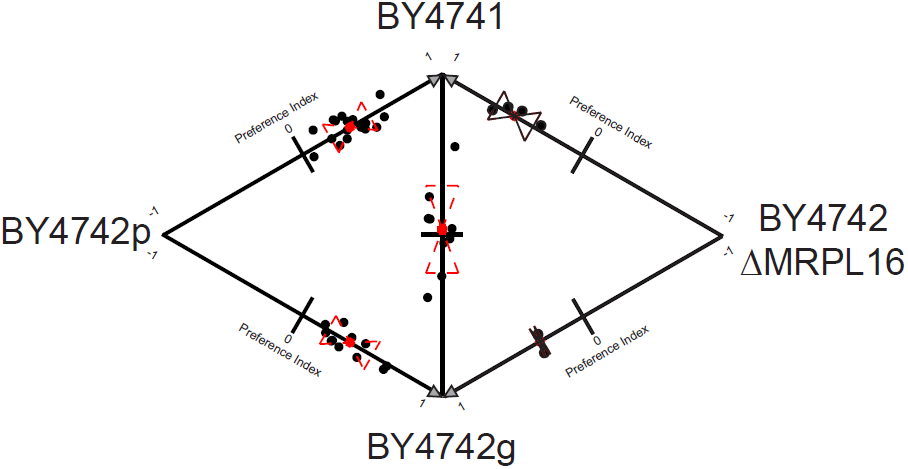
Mitochondrial status of *S. cerevisiae* affects *D. melanogaster* preference. Pairwise behavioral results between BY4741 and various BY4742 strains, each with a different mitochondrial status confirm role of mitochondria in *D. melanogaster* preference. BY4742p is the original isolate, BY4742g is an independent BY4742 strain with functional mitochondria, and BY4742 ΔMRPL16 is a strain from the deletion collection known to genetically disrupt mitochondrial function. Each black dot represents the preference index for each trial (see Figure 1). Red dashed lines represent the two-tailed standard deviation and the red “+” represents the mean of all trials.

### BY4741 and BY4742p strains produce different volatile bouquets, influenced by their mitochondrial status

In the choice assay used here, *D. melanogaster* are selecting between different yeast strains based primarily on how they smell. Thus, the absence of functional mitochondria must be affecting the volatile compounds produced by the yeast strains. We used GC-MS to characterize and quantify the volatile compounds associated with both BY4741 and BY4742p grown on YPD plates, and, as expected, found significant differences (Figure 4A).

**Figure 4.**
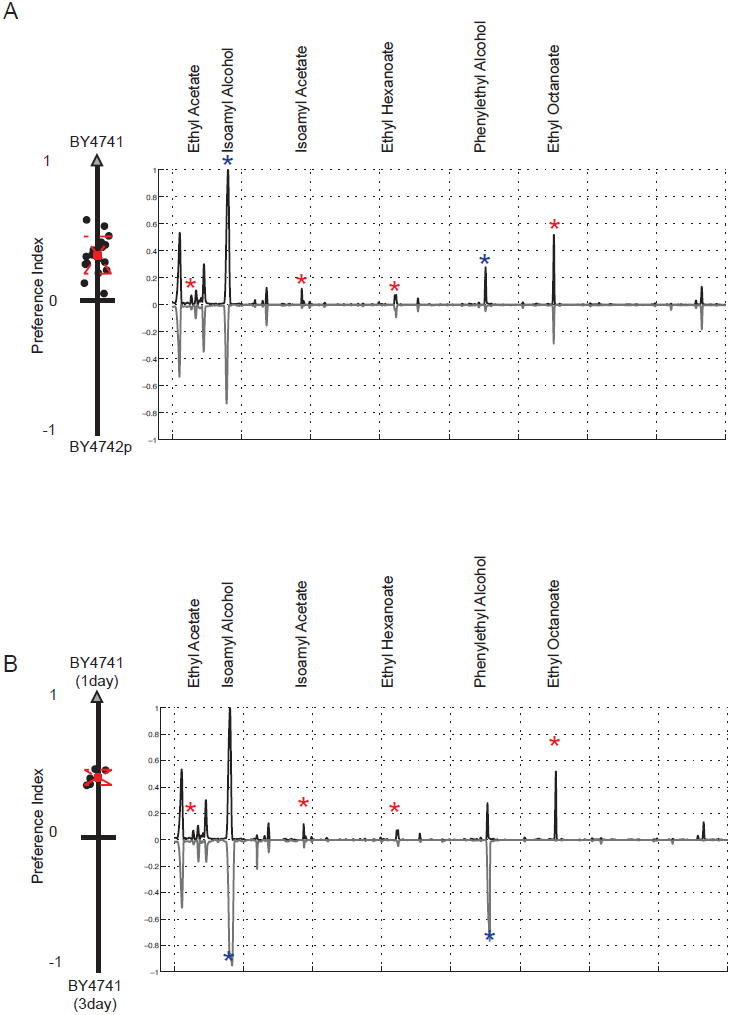
GC-MS identifies volatile compounds that likely underlie *D. melanogaster* preference for *S. cerevisiae* strains with functional mitochondria. A) To evaluate the effect of mitochondrial status on volatile compound production, we performed GC-MS on one day old BY4741 and BY4742p cultures. The average total ion chromatograms (TIC) for BY4741 is plotted on the positive y-axis, while the TIC for BY4742p is plotted on the negative y-axis. The preference data from Figure 3 for these two strains is shown to the left of the panel. Peaks marked with stars represent compounds at higher abundance in BY4741 and represent potential attractants. Compound names are given for all starred peaks. B) To evaluate if the differences were due to the respiration defect in BY4742p, we used GC-MS to compare the volatile profiles of one-day-old BY4741 cultures and respiration-dominated, three day old BY4741 cultures; *D. melanogaster* exhibits a strong preference for the one-day-old cultures. The one day BY4741 TIC is plotted on the positive y-axis, and the three-day-old BY4741 TIC on the negative y-axis. One day BY4741 cultures produce higher levels of a subset of the attractants (red stars), while the others grow in the three-day cultures (blue stars).

To vary mitochondrial engagement, we next examined the volatile profiles of BY4741 and BY4742p in liquid YPD cultures under both anaerobic and aerobic conditions. The anaerobic and aerobic BY4742p cultures produced nearly identical chromatograms (Figure S2). In contrast, the anaerobic and aerobic BY4741 cultures differed, with the anaerobic BY4741 culture resembling the BY4742p culture, and the aerobic BY4741 culture showing a marked increase in the same set of compounds found at higher levels in BY4741 compared to BY4742p plates (Figure S2; see also Figure S3 which shows a similar comparison for one of the sporulated strains with identical genotype to BY4741). These potential attractants include: ethyl acetate, isoamyl alcohol, isoamyl acetate, ethyl hexanoate, phenylethyl alcohol, and ethyl octanoate (Table 1).

**Table 1.**
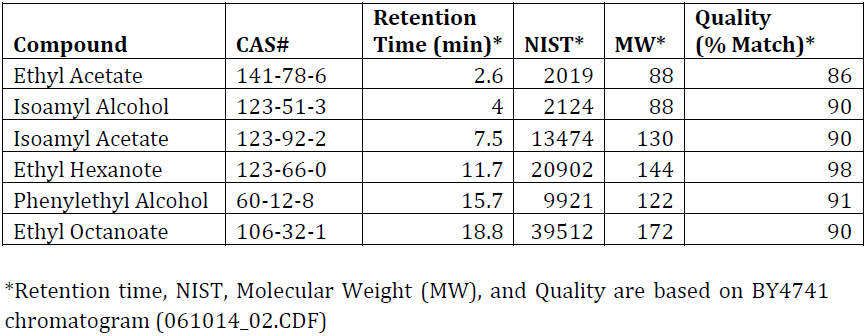
Identified chemical attractants

| **Compound** | **CAS#** | **Retention Time (min)\*** | **NIST\*** | **MW\*** | **Quality (% Match)\*** |
| --- | --- | --- | --- | --- | --- |
| Ethyl Acetate | 141-78-6 | 2.6 | 2019 | 88 | 86 |
| Isoamyl Alcohol | 123-51-3 | 4 | 2124 | 88 | 90 |
| Isoamyl Acetate | 123-92-2 | 7.5 | 13474 | 130 | 90 |
| Ethyl Hexanoate | 123-66-0 | 11.7 | 20902 | 144 | 98 |
| Phenylethyl Alcohol | 60-12-8 | 15.7 | 9921 | 122 | 91 |
| Ethyl Octanoate | 106-32-1 | 18.8 | 39512 | 172 | 90 |

\*Retention time, NIST, Molecular Weight (MW), and Quality are based on BY4741 chromatogram (061014\_02.CDF)

The affect of mitochondria on the volatile profile of a strain is, at least superficially, an unexpected result. *S. cerevisiae* exhibit a phenomenon known as the Crabtree effect, in which they preferentially ferment glucose to ethanol, even in the presence of oxygen [36, 37]. Thus, we would not expect the absence of a functional mitochondria to have a significant effect until available glucose has been depleted and the yeast are forced to respire. However, our data clearly indicate that the mitochondria are having a significant phenotypic effect during both fermentation and respiration.

One possibility is that there is a low level of respiration in cultures that are primarily fermenting. If respiration alone is driving attraction, then we would expect older cultures, which should have higher rates of respiration, to be more attractive. However, we found exactly the opposite: younger cultures of BY4741 are more attractive than older cultures, suggesting that attractiveness is not entirely respiration dependent (Figure 4B).

When we compared the volatile compounds produced by the older and younger cultures using GC-MS we found that several of the potential attractants (isoamyl alcohol, phenylethyl alcohol) continued to grow in concentration in the older, respiring cultures, while the rest (isoamyl acetate, and the ethyl esters) disappeared (Figure 4B). These results suggest that individual metabolites vary in levels of attractiveness. We classified the esters: ethyl acetate, isoamyl acetate, ethyl hexanoate, and ethyl octanoate, which are highest in the younger plates, as primary attractants. We classified isoamyl alcohol and phenylethyl alcohol as secondary attractants, as our results suggest these compounds are not as attractive as the esters.

Overall, these observations suggested that high levels of esters produced by BY4741 during our early time point are primarily driving attraction, and that they are not strictly associated with respiration. Instead, they appear to arise from a separate mitochondria-linked process.

### EEB1 Δ strain affects ethyl ester production and attraction

To further investigate ethyl esters as attractants, we next tested the EEB1Δ strain from the BY4741 knockout collection (*MATa;his3;leu2;met15;ura3;eeb1*). Both EEB1 and its paralog EHT1 have been implicated in medium chain ethyl ester production, with most of the function attributed to EEB1 [38, 39]. We found the EEB1Δ strain to be less attractive than BY4741 in our behavioral assay when strains were grown for the standard incubation time (Figure 5). GC-MS analysis confirmed a decrease in the production of both isoamyl acetate and ethyl esters in the EEB1Δ strain, further implicating these compounds as important attractants in these cultures (Figure 5).

**Figure 5.**
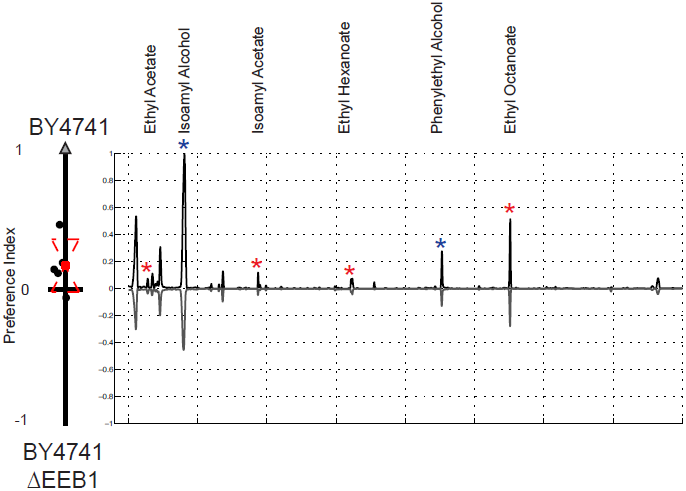
Deletion of EEB1 decreases ethyl ester production and affects *D. melanogaster* preference. Results of preference assay comparing BY4741 and BY4741 ΔEEB1 (left), along with averaged TICs for GC-MS analysis of for BY4741 (positive y-axis) and BY4741 ΔEEB1 (negative y-axis). Both TICs were normalized together so that the maximum value across both experiments was set to 1.

### Nitrogen source affects both the production of identified attractants and fly preference

Although it is not widely appreciated, the mitochondria is required for several metabolic processes not related to respiration [40]. For example, both proline catabolism and branched chain amino acid anabolism have biochemical steps that occur within the mitochondria [41, 42]. We therefore reasoned that varying nitrogen sources – especially those with and without amino acids, or with different distributions of amino acids – might lead to the differential engagement of the mitochondria, different volatile compound production, and different attractiveness to flies.

To test this we grew a prototropic, wild caught *S. cerevisiae* strain, T73, on media containing equivalent carbon source (5% glucose) and varying types and levels of nitrogen (Table 2). We collected GC-MS data for each condition and tested all pairwise comparisons between these media in our behavioral assay. We found that varying the nitrogen source had a dramatic effect on the types and levels of volatile compounds produced by the culture, including the identified attractants from our BY4741 strain (Figure 6). Further, the level of the attractive compounds produced matched with behavior: the culture that produced higher combined levels of attractive compounds in each pairwise comparison was always more attractive to *D. melanogaster* (Figure 6).

**Table 2.** Nitrogen composition of media used in this study

| **Media** | **Ammonium Sulfate (per L)** | **Amino Acids (per L)** | **Glucose (per L)** |
| --- | --- | --- | --- |
| YNB | 5.0g | 0 | 5.0g |
| SC | 0 | 2.0g | 5.0g |
| YPD | >5.0g* | >2.0g* | 5.0g |

\*YPD is a complete media source made by combining yeast extract and peptone as a nutrient and nitrogen source. The exact nitrogen composition of these products is unknown, but each is used in excess in a typical YPD media recipe (represented above as >5.0g/L and >2.0g/L)

**Figure 6.**
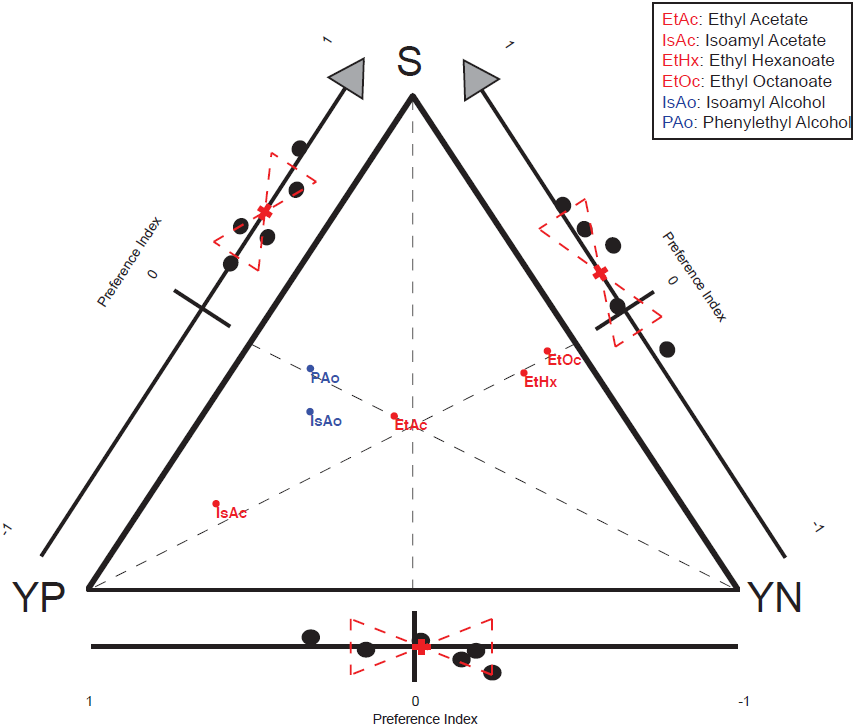
Nitrogen source affects levels of identified attractants and fly behavioral preference in a prototrophic *S. cerevisiae* strain. We performed pairwise preference assays and GC-MS on a prototrophic strain of *S. cerevisiae* (T73) grown on yeast extract, peptone, dextrose media (YPD), yeast nitrogen base media (YNB) and synthetic complete media (SC) which vary in types and levels of nitrogen. The pairwise *D. melanogaster* behavioral results have been added to each side of the triangle representing the three pairwise comparisons. GC-MS data is plotted using triangular Barycentric coordinates (the point for a given compound is placed at the center of mass if the vertices are assigned a mass corresponding to the GC-MS signal for that compound in the corresponding sample). The variation in each attractant for every pairwise comparison can be read by looking at how the compounds segregate across the midpoint of the relevant side (drawn on figure as dashed line for emphasis). Primary attractants are plotted in red and secondary attractants are plotted in blue. Attractants, in combination, track with observed behavioral preferences, but no single attractant can explain all behavioral results.

Interestingly, across all pairwise comparisons, the most attractive medium tested was the synthetic complete medium (SC), which most closely matched the nutrient composition of a fruit, the natural substrate of *S. cerevisiae*. Like fruit, SC media contained a mixed pool of amino acids as a nitrogen source and high levels of sugar (5% glucose) (Table 2). This nutritional composition makes sense for ester production. The high levels of sugar allow for uninhibited fermentation, while the amino acid based nitrogen supply can be more efficiently consumed when the mitochondria are functioning. Ethyl esters are formed from the fermentation byproducts, ethanol and aliphatic acids, largely via EEB1 activity. The reaction requires ATP, which will be far more abundant in fermenting cells with functional mitochondria [38, 39]. We propose that *D. melanogaster* have evolved to sense these esters specifically and use them as a signal of *S. cerevisiae* fermenting a sugar-rich fruit substrate, an ideal oviposition site for *D. melanogaster* females.

### The prototrophic strain, T73, grown on a natural fruit substrate produces the highest levels of identified attractants

Based on the results of our nitrogen experiments, we collected GC-MS profiles of T73 grown on pureed grape agar. The GC-MS profiles for these cultures were dominated by our identified attractants. These cultures contained the highest level of ethyl esters and isoamyl acetate seen in this study (Figure 7). These grape-based fermentations were extremely attractive to *D. melanogaster* and reliably attracted more flies than the most attractive media from previous comparisons (5% SC) (Figure 7).

**Figure 7.**
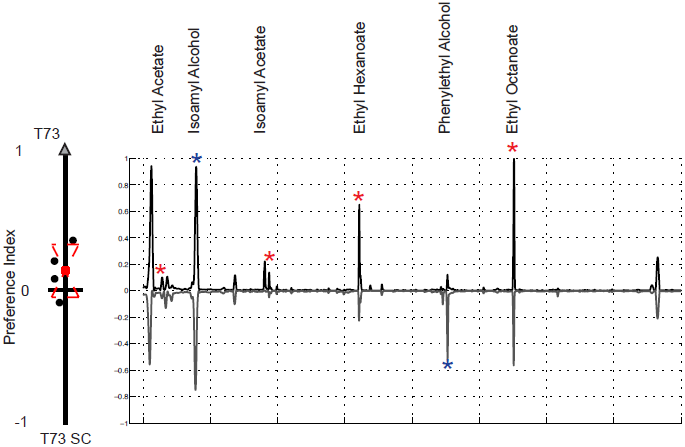
Primary attractants are detected at extremely high levels when T73 is grown on its natural substrate, grape. Behavioral results comparing the prototrophic strain T73 grown on grape media and synthetic complete media (SC) (left). Average TIC for GC-MS data of T73 grown on grape media (positive y-axis) and on synthetic complete media (SC) (negative y-axis). Both TICs were normalized together so that the maximum value across both experiments was set to one.

## Discussion

*S. cerevisiae* is a versatile microbe that can survive and propagate on many nutritional sources. Different combinations of substrate and genotype impose unique metabolic demands on the yeast, leading to the production of distinct volatile profiles. These characteristic volatile fingerprints can, in turn, signal to other organisms, such as *D. melanogaster*, information about the substrate, and, potentially, its suitability for their purposes. In this work, we have identified a piece of the volatile vocabulary that helps bring together *S. cerevisiae* and *D. melanogaster* in nature.

Our research shows that mitochondria affect *S. cerevisiae* metabolism even during periods of fermentation, and that these metabolic alterations dramatically influence the volatile small molecules produced and released by *S. cerevisiae*. In this way, *S. cerevisiae* cells with and without functional mitochondria, grown on the same substrate, produce different volatile profiles. We have shown that *D. melanogaster* are able to sense the differences in the volatiles produced by these strains and prefer the volatile “fingerprint” produced by yeast strains with mitochondria. Using GC-MS, we identified the molecular compounds produced at higher levels in the attractive cultures and tagged these compounds as mitochondrially-associated attractants.

We also found that varying the type of nitrogen provided to a culture changes the levels and varieties of volatiles it produces, including our mitochondrially-associated attractants. This makes sense, as metabolism of many nitrogen sources are known to be associated with mitochondria. We found the highest levels of our attractants when the wild-caught *S. cerevisiae* strain was grown on a media substrate with a nutritional composition similar to that of fruit. Like fruit, the media contained high levels of fermentable sugars and a mixture of amino acids as a nitrogen source – a nutritional scenario lending towards a combination of fermentation and mitochondrial activity.

Together these results suggest that *D. melanogaster's* preference towards mitochondrially-associated metabolites may have an ecological basis. We hypothesize that *D. melanogaster* have tuned their senses towards these mitochondria-associated attractants, as these compounds are produced when *S. cerevisiae* are actively fermenting a fruit, where the combination of yeast and ethanol produce an ideal environment for *D. melanogaster* larvae to develop and remain protected from other microbial threats.

## Methods

### Behavioral setup

A custom-­crafted behavior arena and assay protocol was developed for these experiments. Five behavior assays were conducted in parallel. Twenty-two hours (4pm) prior to starting a trial, the specific yeast strains to be tested were streaked from stock plates (stored at 4C and re-streaked from freezer stock (-80C) every two weeks) onto freshly prepared 60 × 15 mm Petri dish plates (see ‘Media Preparation’ for details). Plates were then transferred to 30C incubator and grown overnight. Twenty-two hours later (2pm, following day) plates were removed from incubator and behavioral traps were assembled for the assay.

To construct the traps, the cultures were topped with a custom-designed and 3D printed lid (uPrint Plus), which was then secured into place with Parafilm. The custom lid allowed for the attachment of a 50 mL Falcon tube (bottom sawed off to create a long, open tube) above the plate. A small piece of metal mesh was secured between the adapter and falcon tube to allow for the volatiles emitted from the culture to dissipate up through the Falcon tube, while physically separating the plate from the Falcon tube chamber. The Falcon tube chamber was finally topped with a funnel fashioned out of Whatman Grade 1 Filter paper (15.0cm diameter). The funnel was secured in place by a small piece of lab tape (Figure S4).

The constructed traps were next transferred into the behavior chambers: 24” high, 12” diameter clear, open-ended, acrylic cylinders (TAP plastics) fitted with closable sleeve that allowed for placement and removal of both the traps and flies (Figure S4). Behavior chambers were kept in a large room with over-head, diffuse lighting that remained on for the entirety of the trial. The chambers were vacuumed and cleaned with water and ethanol before each trail. Four traps were placed in each cleaned arena, two for each yeast strain under analysis. Each strain pair was tested in three different orientations: 2X2 (with the same strain next to each other) in each orientation and a diamond orientation (with the same stain across from the other). One hundred and twenty 4-10 day old *D. melanogaster* were added into the behavioral chamber between 3 and 3:30 pm (see “Fly Husbandry and Handling” section for details). The traps were removed from the chambers 18h later, between 10 and 10:30 am. Before removal the orientation of each trap in each arena was recorded. Flies in each trap were then sexed, counted, and recorded. These raw counts were used to determine the preference for each strain in each arena. The raw data can be found in Spreadsheet 1.

### Calculation of Fly Preference (Preference Index)

To quantify and compare independent behavioral trials, we used a simple preference index:

Every behavioral trial contained four traps. Two baited with yeast strain “A” and two baited with strain “B”. One hundred and twenty 4-10 day old *D. melanogaster* were added into each behavioral trial:

A = Total *D. melanogaster* caught in both traps baited yeast strain “A” in the trial

B = Total *D. melanogaster* caught in both traps baited yeast strain “B” in the trial

**Preference Index (PI) = A**-**B/(A + B)/2**

### Fly Husbandry and Handling

*D. melanogaster* line Raleigh 437 was obtained from Bloomington Stock Center for use in these experiments. All *D. melanogaster* used in behavioral assays were raised using the same regimen. A “time course” for each line was maintained for the duration of the experiments. The time course consisted of pushing 3 vials of 25 flies into fresh media (CSM formula) topped with dried yeast pellets (Red Star) daily. The 25 flies were refreshed with 4-day-old flies from a stock population every 10 days. The time course of vials was kept at 18C and the eggs laid in each vial were allowed to develop (∼18 day generation time). After ∼18 days, each seeded vial was checked daily for newly eclosed flies. All flies hatching off of on a given day were transferred into fresh vials and stored in a 25C incubator for at least 4 days to ensure full development and maturation to adulthood. The entire process was repeated daily to maintain each strain.

To prepare flies for a behavioral trial, aged 4-10 day old flies were removed from the 25C incubator and anaesthetized using CO2 and counted into groups of 120. The groups were visually inspected to insure roughly equal numbers of males and females. The flies were transferred into a vial of fresh food and allowed to recover and eat for 2-4 hours before being added into the behavior chamber.

A stock of Raleigh 437 was maintained at 18C and transferred to fresh media monthly.

### Media Preparation

5% YPD for behavioral assays and strain storage was prepared weekly using the following recipe (for 1L): 20g Peptone (BD Bacto Peptone), 10g Yeast Extract (Amresco Yeast Extract, Bacteriological, Ultra Pure Grade), 50g Dextrose (Fisher Scientific Dextrose Anhydrous), 20g Agar (BD Bacto Agar), Water to 1L. Solution was mixed and heated to boiling on a magnetic stir plate. Liquid YPD was prepared using (for 1L): 20g Peptone, 10g Yeast Extract, 50g Dextrose, Water to 1L. The solution was stirred for five minutes and filter sterilized using Nalgene vacuum filtration system (0.22 micron).

The following additional recipes were used in this experiment:

<u>5% SC (for 1L):</u> 1.7g YNB without amino acids and ammonium sulfate (BD Difco), 2.0g SC Amino acid mixture (MP Biomedicals), 50g Dextrose, 20g Agar, Water to 1L.

<u>5%YNB (for 1L):</u> 6.7g YNB without amino acids (BD Difco), 50g Dextrose, 20g Agar, Water to 1L.

<u>Grape Agar (for 1L):</u> 1.7g YNB without amino acids and ammonium sulfate, 1L organic grapes (v/v) (thoroughly washed and pureed in food processor), 20g Agar, Water to 1L.

### Yeast strains

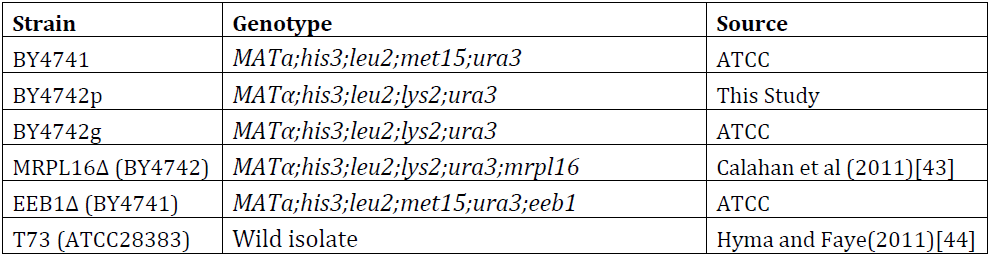

### Tetrad Experiment

Twenty tetrads were obtained using a standard sporulation protocol (Methods in Yeast Genetics, 2005). Briefly, BY4741 and BY4742p were mixed on a YPD agar plate to encourage mating. Diploid colonies were selected by replica plating onto a selective media (Met(-);Lys(-)) transferred to liquid “YPA” media for 1 day, spun down and resuspended in “Minimal Spo” for at least 3 days. Ascii were dissolved and tetrads dissected. After incubation, tetrads were genotyped by replica plating onto selective media for MAT, LYS, and MET loci. Genotyped tetrads were grown in YPD and prepared as freezer stocks using standard protocol (Methods in Yeast Genetics, 2005). Freezer stocks of each stain were used to start cultures for all behavior experiments.

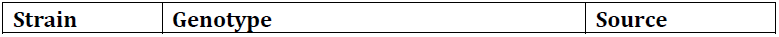
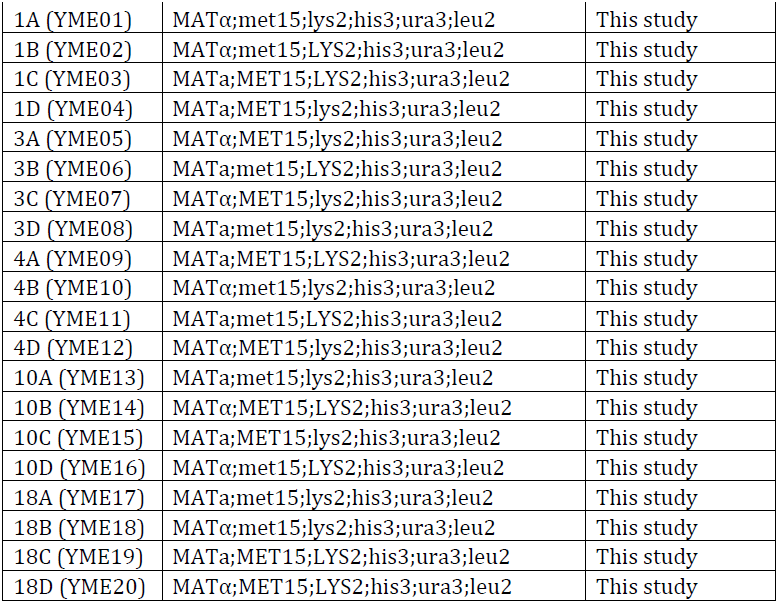

### GC-MS protocol

Sample analysis was performed on Agilent 7890A/5975C GC-MS equipped with a HP-5MS (30m x 0.25mm, i.d., 0.25micrometers film thickness) column.

To sample the headspace of a plate culture, a conditioned, Twister stir bar (10 mm in length, 0.5 mm film thickness, 24microliters polydimethylsiloxane volume) was suspended from the lid of the Petri dish using rare earth magnets for 40 minutes at room temperature. The Twister was then dried using a Kimwipe, placed in a Gerstel thermal desorption sample tube, topped with a transport adapter, and loaded into sampling tray.

Automated sampling and analysis was performed using the Gerstel MPS system and MAESTRO integrated into Chemstation software.

Samples were thermally desorbed using the Gerstel Thermal Desorption Unit (TDU), followed by injection into the column with a Gerstel Cooled Injection System (CIS-4). Temperature program for desorption was 30C (.40 min), then 60C/min to 250C(hold for 5 mins). Temperature of the transfer line was set at 275C. The injection was cooled with liquid nitrogen to -100C with a hold time (.60min). The injector inlet was operated in the Solvent Vent mode, with a vent pressure of 9.1473 psi, a vent flow of 30mls/min, and a purge flow of 6mls/min (for .01 min).

The following GC-MS method parameters was used for each sample: The GC temperature program was 40C (2 mins), then 4C/min to 140C (hold 0 mins), then 15C/min to 195C (hold for 0 min). The carrier gas head pressure was 9.1473 psi for a flow rate of 1.2 mL/min. The GC was operated in the constant flow mode. The MS was operated in EI mode with the electron voltage set at autotune values. The detector was set to scan from 30 to 300amu at a threshold of 150 with 3 samples for a scanning rate of 2.69 scans/second. The MSD transfer line temperature was set at 280C. The ion source and quadrupole temperatures were set at 230C and 150C, respectively.

GC-MS data files were visually inspected using Chemstation. TIC peaks were identified using the NIST O8 database via Chemstation. Datafiles were transferred, parsed, and analyzed using custom written Matlab scripts (see “Scripts Supplement”).

### Aerobic and anaerobic yeast cultures

Pairs of 100mL 5% YPD cultures were inoculated with BY4741 or BY4742p cells (stored on a solid 5% YPD plate at 4C) and topped with either a loose foil cap or fermentation bung (Ferm-Rite) to create aerobic and anaerobic growth conditions for each strain. Cultures were grown for 21h shaking at room temperature. After this growth period, cultures were removed from shaker, and an O.D. reading was taken to ensure similar cell densities across each aerobic and anaerobic culture pair. 50mL of each culture was transferred to a .22 micron vacuum filter (MilliQ Steri-Flip) to remove yeast from the culture medium. 20mL of the culture medium was then transferred into 20mL amber glass vials (Sigma), a conditioned Twister stir bar (Gerstel) was added, and the vial was capped with a silica-lined screw cap (Sigma). The Twister stir bars were spun in the medium at 300rpm for 40min, rinsed with MilliQ water, dried with a Kimwipe, and transferred onto the GC-MS for analysis (see “GC-MS protocol” for details).

### Glycerol plate growth

BY4741 and BY472p and tetrads representing each genotype were streaked from 5% YPD stock plates onto 5% glycerol plates. Plates were incubated at 30C for 48h and visually inspected for growth.

## Acknowledgments

KMS would like to thank Meru Sadhu for help with tetrad dissection experiment, Tommy Kaplan and Peter Combs for advice on Matlab scripts, Steven Kuntz for advice on figures, and Kyle Barrett for support. MBE would like to thank Barbara Dunn, whose fascination with yeast ecology in the Botstein lab in the 1990s inspired these experiments.

---

**Supplemental Figure 1.**
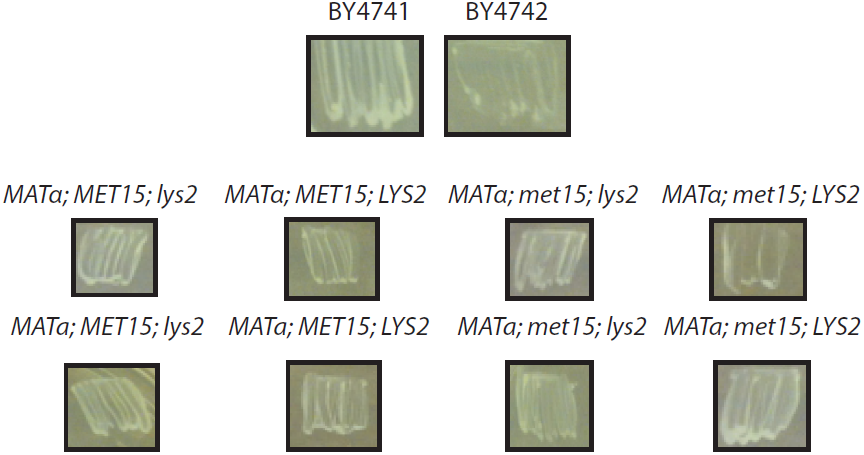
Confirmation of mitochondrial deficiency in BY4742p. BY4741, BY4742p, and tetrads (labeled by genotype) grown on glycerol to check for ability of each strain to respire. All strains except BY4742p showed visible growth.

**Supplemental Figure 2.**
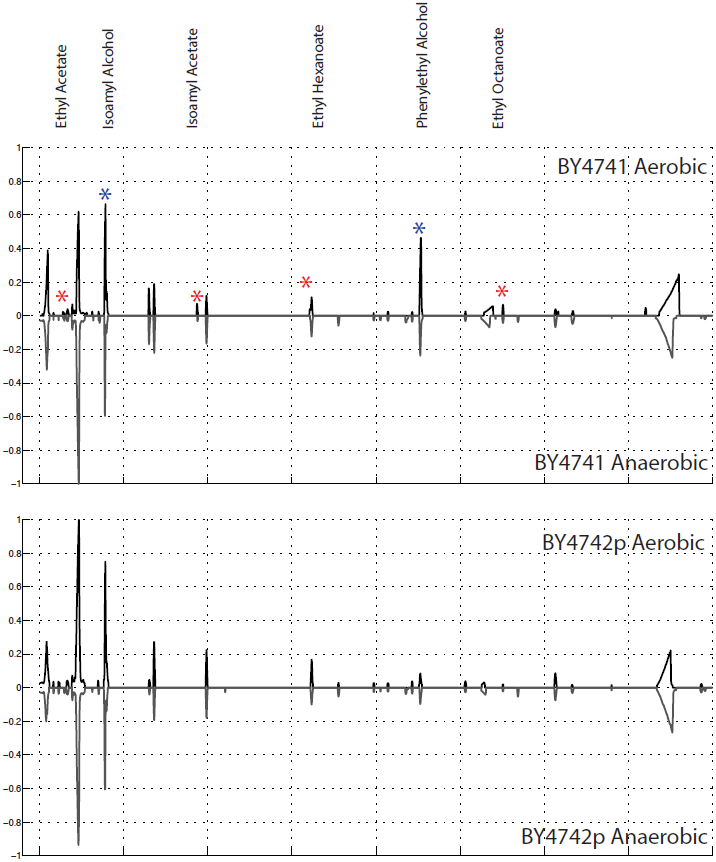
Comparison of GC-MS profiles of aerobically and anaerobically grown BY4741 and BY4742p cultures. Averaged TICs for: A) BY4741 grown both aerobically and anaerobically and B) BY4742p grown aerobically and anaerobically in 5% YPD. TICs in each panel were normalized together so that the maximum value across both experiments was set to one. In each comparison, aerobic cultures are plotted on the positive y-axis and the anaerobic cultures plotted on the negative y-axis.

**Supplemental Figure 3.**
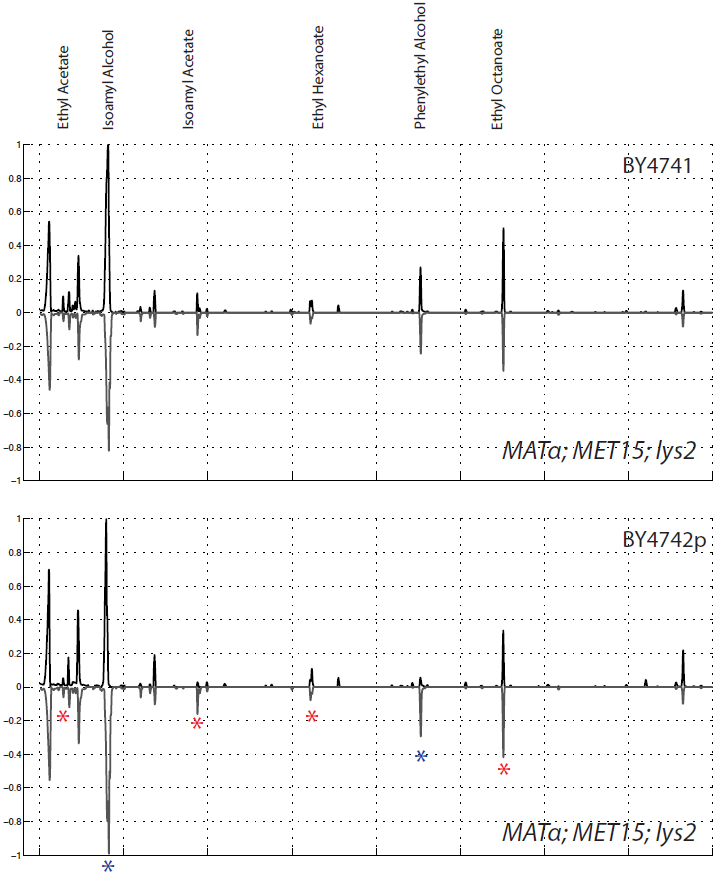
Tetrad produces GC-MS profile similar to BY4741 at identified attractants. Averaged TICs for BY4741, BY4742p and a selected tetrad (*MATα;MET15;lys2*). BY4741 and BY4742p are plotted on the positive y-axis and the tetrad on the negative y-axis in each panel. TICs in each panel were normalized together so that the maximum value across both experiments was set to one.

**Supplemental Figure 4.**
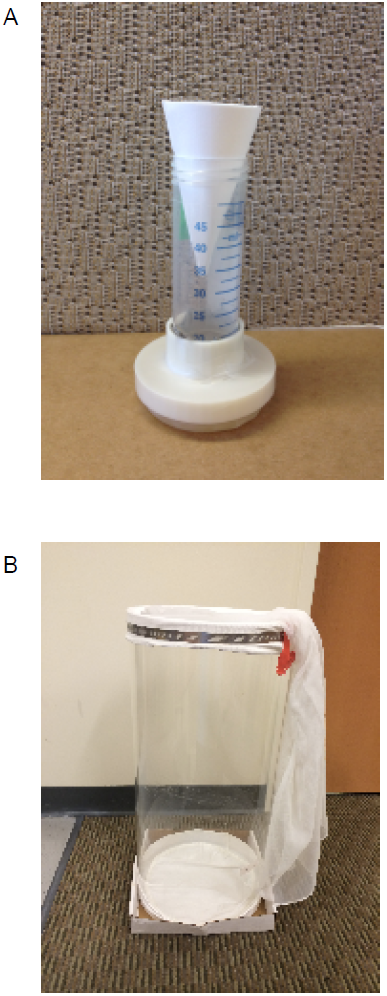
Images of the behavioral trap (A) and behavioral arena (B) explained in “Behavioral setup” section of Methods.

